# Environment-dependent epistasis shapes adaptation to fluctuating resources

**DOI:** 10.64898/2026.07.31.742109

**Authors:** Ashlynn G. Bruder, Nathan Fiedeldey, Sarah B. Worthan, Zeer Cen, Megan G. Behringer

## Abstract

Microbial populations frequently experience fluctuations in resource availability, yet how these fluctuations reshape genetic interactions during adaptation remains unclear. Here, we identify the rapid evolution of latent antimicrobial susceptibility in *Escherichia coli* populations experimentally evolved under repeated feast-famine cycles. Longitudinal population genomics revealed recurrent mutations in the transport genes *acrB* and *ompF*, and reconstruction of these mutations demonstrated environment-dependent fitness effects characterized by positive epistasis after 1 day of culture and reciprocal sign epistasis after 10 days. Together, our results support a two-step adaptive trajectory in which mutations affecting efflux are followed by changes in porin permeability, generating genotypes that are jointly beneficial across feast and famine. More broadly, our findings demonstrate how fluctuating environments reshape adaptive genetic interactions, producing historically contingent evolutionary trajectories and latent phenotypes.

**Article Summary:** How fluctuating environments shape genetic interactions and evolutionary trajectories remains a central question in evolutionary genetics. We identified latent antimicrobial susceptibility in *Escherichia coli* populations evolved under repeated feast-famine cycles and reconstructed evolved mutations to resolve the genotype-to-phenotype map underlying this trait. The responsible mutations arose in key transport genes, exhibited environmental-dependent epistasis, and increased fitness across both feast and famine conditions. Our findings provide experimental evidence that fluctuating environments can select for interacting mutations that optimize fitness across a dynamic adaptive seascape.

## Introduction

Microbial populations experience fluctuating environmental conditions in nearly all habitats, including soil, aquatic systems, host-associated environments, and laboratory culture (J. Nguyen et al., 2020; Savageau, 1983; Zhang & Rubin, 2013). Among these fluctuations, changes in nutrient availability are especially pervasive and often occur as repeated cycles of resource abundance (“feast”) followed by periods of resource limitation (“famine”). Because the physiological requirements for rapid growth differ from those needed for long-term survival, adaptation to fluctuating resource conditions must balance rapid and efficient growth during feast but also maintain the population during famine (Aertsen & Michiels, 2004; Niimi et al., 2026). Consequently, the temporal structure of nutrient availability can strongly influence evolutionary trajectories by altering the selective pressures experienced throughout the microbial growth cycle (Kussell & Leibler, 2005; J. Nguyen et al., 2021).

Studies of adaptation to resource limitation have revealed tradeoffs between traits that enhance growth and those that promote survival (Finkel, 2006; Manhart et al., 2018; Vasi & Lenski, 1999; Zhu & Dai, 2024). These tradeoffs arise from bioenergetic constraints on resource allocation, thermodynamic constraints of metabolic pathways and enzymatic activity, and constraints on global regulation of gene expression (Biselli et al., 2020; Caetano et al., 2021; Ferenci, 2005; Hui et al., 2015; Mori et al., 2017; Schink et al., 2022). Under fluctuating conditions, adaptation may also promote mutations whose effects depend on ecological context and genetic background (Cvijović et al., 2015; Hall et al., 2019; Karita et al., 2026). These interactions may also generate epistatic genetic architectures based on mutations with pleiotropic effects: improving performance across multiple phases of the extended microbial growth curve while also producing indirect effects on traits not under selection (Anderson et al., 2021; Baier et al., 2023; Behringer et al., 2024; de Vos et al., 2013). These indirect outcomes, often referred to as latent phenotypes, arise as byproducts of adaptation and may influence fitness in alternative environments (Payne & Wagner, 2014).

Understanding the genetic basis of these latent phenotypes is central to explaining how adaptation in one environment influences performance in another. Antimicrobial susceptibility represents a particularly important latent phenotype, as it can be strongly influenced by cellular processes that are not directly related to antibiotic exposure (Knöppel et al., 2017). These effects are often mediated by global physiological changes, such as the stringent response, which reprogram cellular metabolism, growth, and envelope composition under nutrient stress (Corrigan et al., 2016; Laubacher & Ades, 2008; D. Nguyen et al., 2011). Given that adaptation to antibiotic stress can influence bacterial growth parameters (Fitzsimmons et al., 2010), genetic changes that improve fitness in fluctuating environments may also modify susceptibility to antibiotics as an indirect consequence.

In this study, we investigate how adaptation to fluctuating nutrient environments shapes antimicrobial susceptibility as a latent phenotype. We focus on the Long-Term Repeated Starvation (LTRS) experiment, in which populations of *Escherichia coli* have evolved for more than 1,900 days under 1-, 10-, or 100-day cycles of feast and famine (Behringer et al., 2022, 2024; Ho et al., 2021) (**Figure 1**). These populations provide a unique opportunity to examine how differences in the timing and intensity of nutrient limitation influence evolutionary outcomes. We initially predicted that adaptation to repeated starvation would promote generalized stress resistance and increased antimicrobial resistance as a latent phenotype. To investigate this relationship, we quantified antimicrobial susceptibility in evolved clones, used longitudinal genomic data to identify parallel mutations associated with susceptibility changes, and reconstructed candidate mutations in the ancestral background to measure their effects on antimicrobial susceptibility and fitness. By integrating longitudinal population genomics, allele reconstruction, and environment-specific fitness assays, this work demonstrates how adaptation to fluctuating resource environments generates environment-dependent epistatic interactions that shape both selected traits and latent phenotypes.

**Figure 1:**
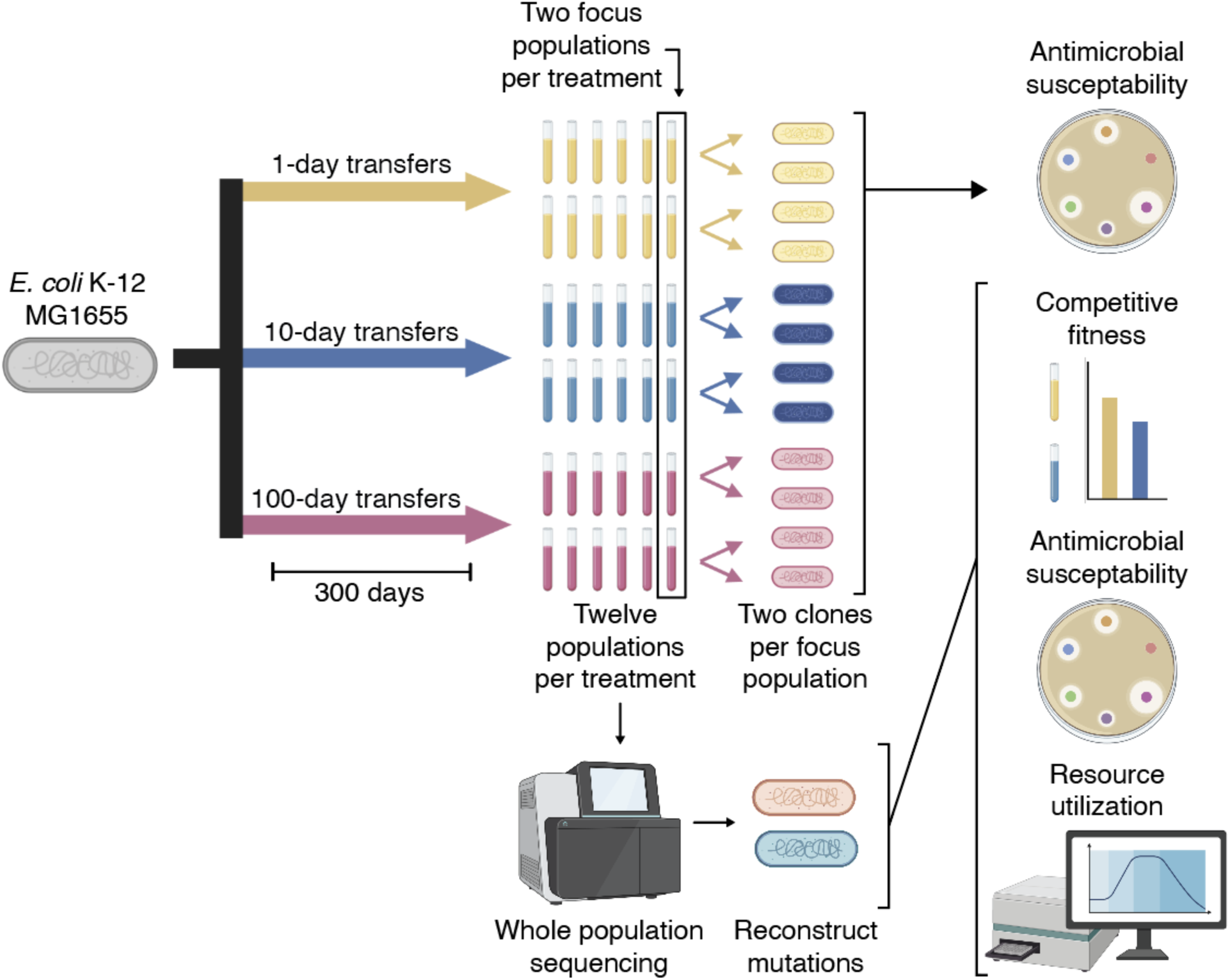
Visual description of study design and methods. Thirty-six populations were each initiated by an independent colony of the Escherichia coli str. K12 substr. MG1655 PMF2 ancestral strain. Populations were split into three groups of 12 and evolved experimentally under 1-, 10-, or 100-day transfer regimes. After 300 days of evolution, clones were isolated from two focal populations for each treatment and screened for antimicrobial susceptibility. Based on these results, we identified candidate mutations from the longitudinal whole-population metagenomic sequencing. After reconstructing these mutations in a clean ancestral background, we assessed fitness, antimicrobial susceptibility, and resource utilization.

## Materials and Methods

### Cultivation of evolved populations and isolation of clones

Experimentally evolved *Escherichia coli* populations used in this study were developed as previously described (Behringer et al., 2022, 2024; Ho et al., 2021). Briefly, 48 populations were evenly distributed across three treatments and propagated in 16 × 100 mm glass culture tubes containing 10 mL of LB Miller broth. Every 1-, 10-, or 100-day interval, we transferred 1 mL of each population into fresh LB medium, resulting in a 1:10 dilution of each population with each transfer, for a total of 900 days. Half of the populations were founded from independent colonies of *E. coli* strain PFM2, a prototrophic derivative of *E. coli* K-12 strain MG1655, while the remaining populations were founded from PFM5, a derivative of PFM2 carrying an in-frame deletion of *mutL* (Lee et al., 2012). Additionally, half of all populations contained a Δ*araBAD56*7 deletion, allowing differentiation of strains by producing red colonies when cultivated on tetrazolium arabinose (TA) agar. All populations were metagenomically sequenced at 100-day intervals to track genomic evolution over time.

At day 300, two focal populations derived from the PFM2, Δ*araBAD56*7 background were selected from each treatment (1-day: P103, P108; 10-day: P403, P408; 100-day: P503, P508). These populations were selected 1) because they carried the Δ*araBAD56*7 neutral marker allowing for competitive fitness assessment against the PFM2 experimental ancestor (hereafter, referred to as WT); and 2) they began experimental evolution as MutL^+^, increasing the likelihood that we can connect genotype to phenotype. To broadly represent the genetic diversity across this study in our phenotypic analyses, we isolated 8 clones from each population for whole-genome sequencing. After sequencing, we mapped the reads to the *E. coli* K-12 MG1655 reference genome (NC_000913.3), identifying the mutations using GATK best practices (SNMs and Indels; https://gatk.broadinstitute.org/hc/en-us ) and GRASPR (structural variants) (Lee et al., 2014). We then selected two distinct clones from each population (12 clones total; **Supplemental Dataset S1**) for antimicrobial susceptibility testing.

### Antimicrobial susceptibility testing

Antimicrobial susceptibility was assessed for each clone, in triplicate, using modified Kirby-Bauer disk diffusion assays (Hudzicki, J., 2009). We selected the Kirby-Bauer method because the diffusion of antimicrobials from an impregnated disc results in a zone of inhibition (ZOI) with a measurable area. Thus, enabling continuous quantitation of antimicrobial resistance in contrast to other methods, such as E-tests or MIC testing in microwell plates. Additionally, by focusing on “first-generation” antimicrobials, we expect to observe the largest phenotypic changes compared to later-generation antimicrobials, which have been engineered for expanded coverage and increased effectiveness. Evolved clones were revived from cryogenic storage by streaking onto LB agar and incubating overnight at 37℃. Individual colonies were then suspended in 600 μL of phosphate-buffered saline (PBS) in 1.5mL microcentrifuge tubes. Cell densities were standardized by measuring absorbance at 600nm with a VWR V-1200 spectrophotometer. Suspensions were adjusted with PBS to an absorbance between 0.10 - 0.13, corresponding approximately to a ∼0.5 McFarland standard. 350 μL of each suspension was spread onto 150 mm Mueller-Hinton agar (MHA) plates (Difco, 275730). Plates were pre-marked into six equal sections to prevent overlap of zones of inhibition (ZOIs). Antimicrobial disks (BD BBL Sensi-Disc) were then applied to each plate, representing six antibiotic classes: erythromycin (ERY 15 μg), nalidixic acid (NAL, 30 μg), penicillin (PEN, 10 U), streptomycin (STR, 10 μg), tetracycline (TET, 30 μg), and trimethoprim–sulfamethoxazole (TMS, 1.25 μg/23.75 μg ). These antibiotics were selected to represent diverse mechanisms of action and cellular targets (Hutchings et al., 2019). Plates were incubated at 37℃ for 20 h, after which they were imaged and analyzed using ImageJ (citation). Each plate image included a 20 mm scale for calibration. The areas (*a*) of both the Zone of Inhibition (ZOI) and the antibiotic disk were measured digitally with ImageJ, and the radius (r) of each was calculated as 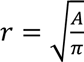 (Schneider et al., 2012). Antimicrobial susceptibility was reported as the net radius *r_net_* = *r_ZOI_* – *r_disc_*.

### Identification of candidate genes contributing to antimicrobial susceptibility

To identify candidate genes potentially contributing to antimicrobial susceptibility phenotypes, we used the following criteria: 1) genes with mutations occurring only in sequenced clones from the 10-day treatment, and not in sequenced clones from the 1-day or 100-day treatments; and 2) genes that were also over-represented for mutations across all of 10-day feast-famine populations in Behringer and Ho, 2022. This resulted in a list of 10 candidate genes (*acrA, acrB, argR, fabF, ompF, oppA, plsX, putP, rpoB,* and *rpsL*). We then performed an enrichment analysis using DAVID v. 2026_1 (https://davidbioinformatics.nih.gov/ ), which returned one significant functional cluster, beta-lactam resistance, containing *acrA*, *acrB*, and *ompF*. Thus, we focused our downstream analyses on these genes (**Supplemental Dataset S2**).

### Reconstruction of key mutations

To investigate the observed changes in antibiotic susceptibility, candidate mutations (*ΔacrB* and *ompF* ^E139A^) were reconstructed in the WT PFM2 ancestral background. This approach allowed us to generate strains containing each mutation individually, as well as in combination. We replaced the WT *ompF* allele with *ompF* ^E139A^ using a CRISPR-FRT-based protocol (Swings et al., 2018). First, a *ΔompF* strain was generated via P1 transduction by moving the *ΔompF846::kan* cassette from the Keio collection strain JW0912 into the PFM2 WT background. The kanamycin cassette was subsequently removed from the *ΔompF* strain by introducing the FLP recombinase expression plasmid, pCP20, and selecting for ampicillin-resistant transformants at 30℃. The plasmid was then cured through nonselective growth at 42℃, and colonies were screened for loss of kanamycin resistance, confirming the *ΔompF846::frt* genotype.

To introduce the *ompF* ^E139A^ allele, the resulting *ΔompF846::frt* strain was then transformed with the pCas plasmid, and transformants were selected at 30℃. Cultures of *ΔompF846::frt* pCas were grown to an OD600 of 0.35, at which point 10 mM arabinose was added to induce expression of the λ red phage proteins. Cells were subsequently prepared as electrocompetent, resuspended in 10% glycerol, and flash frozen in liquid nitrogen for later use. To enable CRISPR-based editing at the FRT site, we generated a guide RNA (gRNA) targeting the sequence 5’-TCCTATTCTCTAGAAAGTAT-3’. This was accomplished using the NEB Q5 site-directed mutagenesis kit (product number: E0554S) to replace the original gRNA sequence present in the pTargetF plasmid (5’-CATCGCCGCAGCGGTTTCAG-3’), resulting in the modified plasmid pTargetF::FRT. A donor DNA template containing the *ompF* ^E139A^ mutation was generated by PCR amplification of ∼500 bp regions upstream and downstream of the mutation from evolved clone number 403-2 (10-day Population A Clone 1; **Supplemental Dataset 1**) using the following primers: forward: 5’-ACGTCTGTGTATTTCTGTGG -3’; reverse: 5’- CAGATCAACATCACCGATACC -3’. Electrocompetent Δ*ompF*846::frt pCas cells were co-transformed with 500 ng of donor DNA and 100 ng of pTargetF::FRT. Transformants were selected on LB agar containing kanamycin (50 µg/mL) and spectinomycin (50 µg/mL) at 30 °C.

The *ΔacrB* mutant strains were generated via P1 transduction by transferring the *ΔacrB747::kan* cassette from the Keio collection strain JW0451 into both the WT PFM2 and PFM2 *ΔaraBAD* backgrounds (Baba et al., 2006). The kanamycin cassette was again removed from the *ΔacrB* strain by introducing the FLP recombinase expression plasmid, pCP20. Lastly, the Δ*acrB/ompF* ^E139A^ double mutant strain was subsequently constructed by introducing the *ompF* ^E139A^ allele into the Δ*acrB/ΔaraBAD* strain using CRISPR-FRT. Final reconstructed strains were verified by PCR and Sanger sequencing, and plasmid loss was confirmed by antibiotic-sensitivity screening, ensuring that any antimicrobial phenotypes were the sole product of allele replacement.

### Assessment of competitive fitness

The fitness of reconstructed mutants was measured relative to the ancestral PFM2 strain using competitive co-culture assays (Worthan et al., 2023). Strains were revived from cryogenic stocks by streaking onto LB agar and incubating at 37 °C for 24 h. To precondition strains to the competition environment, a single colony was inoculated into 10 mL of LB broth in a 16 × 100 mm glass culture tube and incubated at 37 °C with shaking at 180 rpm for 24 h. Cultures were then normalized to equal cell densities based on OD600 measurements using an Epoch2 microplate spectrophotometer. Competition assays were initiated by inoculating 50 µL of the ancestral PFM2 culture and 50 µL of a reconstructed mutant into 10 mL of fresh LB broth. Cultures were vortexed thoroughly, and a 100 µL sample was immediately collected for colony-forming unit (CFU) enumeration (T0). Co-cultures were then incubated at 37 °C with shaking at 180 rpm.

After either 1- or 10-days of competition, cultures were vortexed again, and a 100 µL sample was collected for CFU quantification. At both initial (*T*_-_) and final timepoints (*T*. or *T*_.-_), samples were plated on tetrazolium arabinose (TA) agar. Colony color was used to distinguish strains, with pink colonies corresponding to the PFM2 ancestor and red colonies to the evolved or reconstructed strains. As 10-day competitions span all phases of microbial growth, fitness was quantified using the selection rate 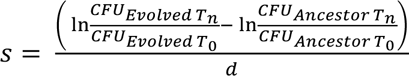 Where *Tn* represents the final time point (day 1 or 10), and *d* is the duration of the competition in days. We then applied the additive epistasis model to assess the existence of epistatic interactions between alleles ε = *S_AB_* – (*S_A_* + *S_B_*) (Phillips, 2008).

To resolve how the reconstructed mutations contributed to fitness under resource depleted conditions, we also performed competitive co-culture in cell-free “spent” or preconditioned media. Preconditioned media was prepared by inoculating a single WT colony into a flask containing 250 mL of LB broth and incubating under shaking conditions (180 rpm) at 37 °C for 24 h. After 24 hours, cells were removed using a 500 mL Vacuum Filter/Storage Bottle System fitted with a 0.2 μm filter (Corning, 431097). Preconditioned media was immediately used after preparation for competitive co-culture assays. A sample of uninoculated preconditioned media was also streaked on to a TA agar plate as a control to ensure that all WT cells were removed.

### Assessment of differential resource utilization

To assess differential resource utilization for the WT and *ompF* ^E139A^ strain, we performed growth assays in Biolog PM1 Phenotype MicroArray plates, in which each well contains a distinct carbon source. Strains were first preconditioned by inoculating a single isolated colony into 16 x 100 mL glass culture tubes containing 10 mL of LB broth and incubated with shaking for 24 h at 37 C. Following incubation, 1 mL of culture was transferred to 1.5 mL microcentrifuge tubes and pelleted by centrifugation at 10,000 rpm for 3 min. Pellets were washed by resuspension in 1 mL of 1x M9 minimal salts, followed by centrifugation under the same conditions. This wash step was repeated five times to remove residual nutrients. After the final wash, cells were resuspended in 1x M9 minimal salts and adjusted to 85% transmittance at 600 nm using a VWR V-1200 spectrophotometer. Each well of the Biolog plate was inoculated with 100 µL of the standardized cell suspension. Plates were incubated at 37 °C in a BioTek Synergy H1 plate reader with continuous orbital shaking. Optical density measurements at 600 nm were recorded every 15 min for 16 h. Each strain was assayed with three biological replicates, and the empirical area under the curve (AUC_E_) was measured using the Growthcurver package v.0.3.1 for R. Resources were removed from the analysis if max(AUC_E_) < 250 across all strains and replicates, resulting in 65 carbon sources for statistical analysis (see **Supplemental Figures S3 and S4**).

### Statistical analysis

All statistical analyses were performed using R Studio v. 2026.05.0 running R v.4.6.0. All t-tests were performed using the compare means function from the ggpubr package v. 0.6.3. Plots were generated using tidyverse v.2.0.0 and ggplot2 v.4.0.3 (Wickham et al., 2019). As the Gen5 software (BioTek) reports time as hh:mm:ss, these values were converted into minutes as decimal values with the lubridate package v.1.9.5. All R code generated for data visualization and statistical analysis is available on the Behringer Lab GitHub (https://github.com/BehringerLab/Environment_Dependent_Epistasis).

## Results

### Antimicrobial susceptibility increases after evolution under 10-day feast–famine cycles

After 300 days of experimental evolution under repeated 1-, 10-, or 100-day feast-famine cycles, we selected two clones from each of two focal populations per treatment (12 clones total) for phenotypic assessment (Behringer et al., 2022, 2022; Ho et al., 2021) (**Figure 1; Supplemental Dataset 1**). We initially hypothesized that adaptation to repeated starvation would produce generalized stress resistance and, consequently, increased antimicrobial resistance as a latent phenotype. To test this, we quantified antimicrobial susceptibility using disc-diffusion assays with six major antimicrobials representing diverse mechanisms of action and cellular targets: erythromycin, nalidixic acid, penicillin, streptomycin, tetracycline, and trimethoprim–sulfamethoxazole (Hutchings et al., 2019) (**Figure 2A**).

**Figure 2:**
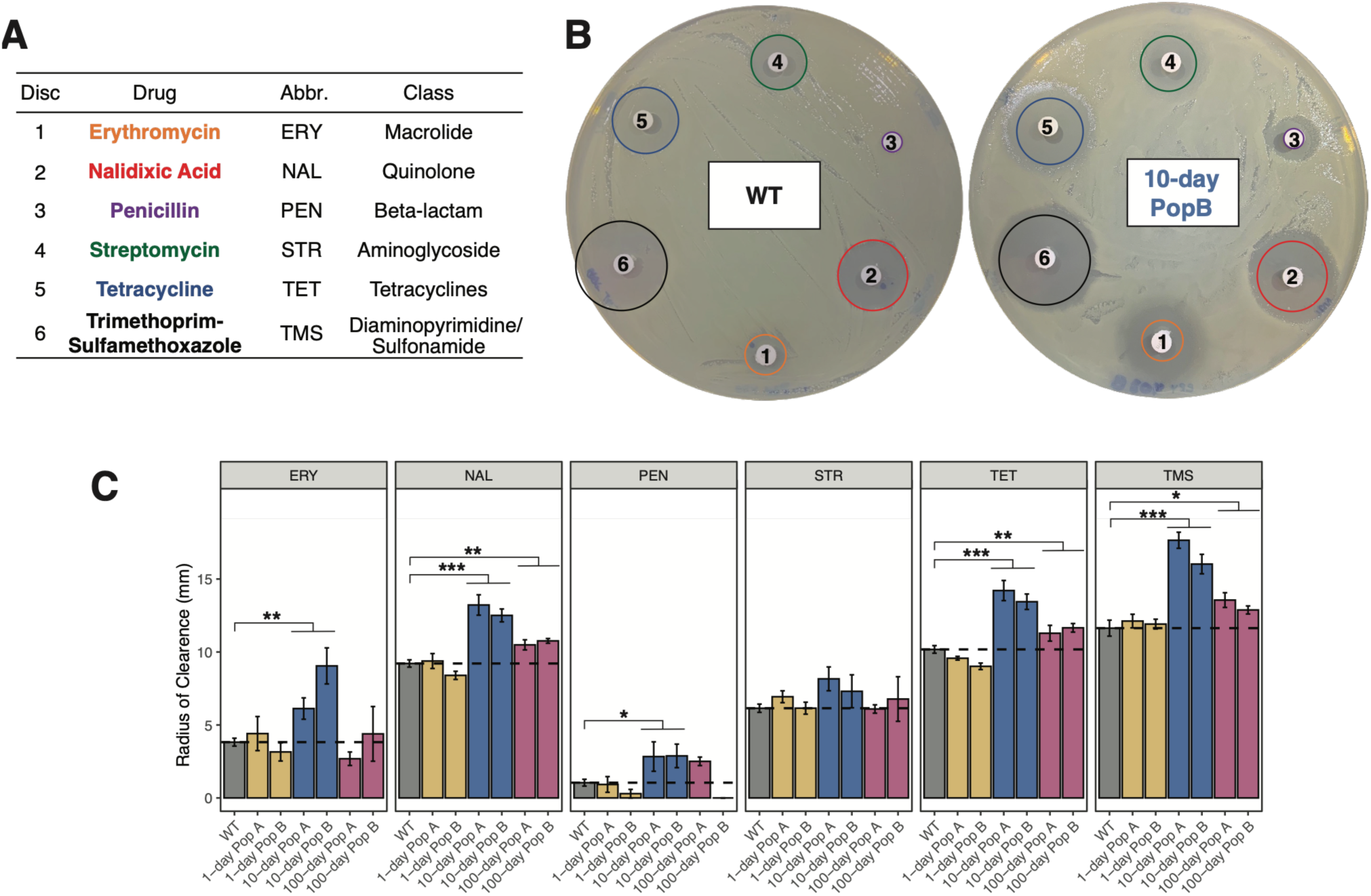
Increased susceptibility to antimicrobials is a unique outcome of evolution under 10-day feast/famine conditions. **A)** Table describing the six antimicrobials used to screen evolved changes in antimicrobial susceptibility. **B)** Side-by-side plate images demonstrating representative changes in the zones of inhibition (ZOI) for each antimicrobial drug in a WT ancestor and a 10-day Population B clone. Numbers identify each antimicrobial disc, and colors represent the ZOI for the WT ancestor projected onto each plate, highlighting the increased ZOIs for the 10-day Population B clone. Numbered discs and colored ZOIs are as follows: 1) Erythromycin (orange), 2) Nalidixic Acid (red), 3) Penicillin (purple), 4) Streptomycin (green), 5) Tetracycline (blue), and 6) Trimethoprim-Sulfamethoxazole (black). **C)** Radii of clearance per antimicrobial averaged across clones from each focus population after 300 days of evolution. Radii of clearance were calculated by measuring the area of the ZOI and the antimicrobial disc, calculating the radius for both, and taking the difference. Black dashed lines indicate the WT ancestor’s radius of clearance for each antimicrobial. Error bars represent +/- s.e.m. *P < 0.05; **P < 0.01; ***P < 0.001.

Contrary to our expectation, evolved clones did not show a general increase in antimicrobial resistance. Instead, we observed a treatment-specific increase in antimicrobial susceptibility, with the strongest effect in clones evolved under 10-day feast-famine cycles (**Figure 2B-C**). Clones from the 10- day feast/famine treatment were significantly more susceptible to five of the six antimicrobials tested. The largest fold-change was observed for penicillin, for which the mean net radius of inhibition increased 2.75-fold relative to the WT ancestor (*P* = 0.046; pairwise t-test). The largest absolute increase was observed for trimethoprim-sulfamethoxazole, with a 5.24 mm increase in mean net radius of inhibition (*P* = 2.8x10^-7^, pairwise t-test; **Table 1**).

**Table 1:** Antimicrobial susceptibility after evolution under different feast-famine regimes.

| Antimicrobial | Ancestor | 1-day |  | 10-day |  | 100-day |  |
| --- | --- | --- | --- | --- | --- | --- | --- |
|  | ROI<br>(mm) | ROI<br>(mm) | p-value<br>(adj.) | ROI<br>(mm) | p-value (adj.) | ROI<br>(mm) | p-value<br>(adj.) |
| ERY | 3.82 | 3.78 | 1 | **7.59** | **0.002^** | 3.54 | 1 |
| NAL | 9.21 | 8.89 | 0.44 | **12.9** | **0.00000099^** | **10.6** | **0.00027^** |
| PEN | 1.04 | 0.608 | 0.55 | **2.86** | **0.046^** | 1.25 | 0.71 |
| STR | 6.15 | 6.54 | 0.71 | 7.73 | 0.14 | 6.44 | 0.72 |
| TET | 10.2 | **9.3** | **0.008\*** | **13.8** | **0.0000021^** | **11.5** | **0.008^** |
| TMS | 11.6 | 12 | 0.54 | **16.8** | **0.00000028^** | **13.2** | **0.036^** |

ROI (mm) = Radius of Inhibition
^ Increased Susceptibility
\* Decreased Susceptibility

In contrast, clones from the 100-day treatment exhibited a more limited susceptibility phenotype, with increased susceptibility observed for three antimicrobials: nalidixic acid, tetracycline, and trimethoprim-sulfamethoxazole. Clones from the 1-day treatment showed little evidence of increased susceptibility and instead exhibited increased resistance to tetracycline (*P* = 0.008, pairwise t-test). Together, these results indicate that adaptation to intermediate feast–famine cycles produced collateral effects on antimicrobial susceptibility, revealing a latent phenotype that was not directly selected during experimental evolution.

### Mutations in acrAB and ompF repeatedly arise in 10-day feast-famine populations

Because clones from the 10-day feast-famine treatment exhibit the greatest increases in antimicrobial susceptibility, we re-examined the longitudinal metagenomic sequencing data from the LTRS populations to identify candidate mutations that could contribute to this latent phenotype (Behringer et al., 2022, 2024; Ho et al., 2021). We identified two repeatedly mutated loci with known roles in antimicrobial susceptibility: the *acrAB* efflux operon and the outer membrane porin, *ompF*. Mutations in these loci were common among 10-day feast-famine populations, with 7 out of 16 populations evolving one or more mutations in *acrAB,* and 9 out of 16 populations evolving one or more mutations in *ompF* within the first 400 days of evolution (**Figure 3A**). By day 900, all 16 populations had evolved mutations in both *acrB* and *ompF*. These repeated mutations suggest that alteration of efflux and outer membrane permeability may be a common feature of adaptation to 10-day feast-famine cycles.

**Figure 3:**
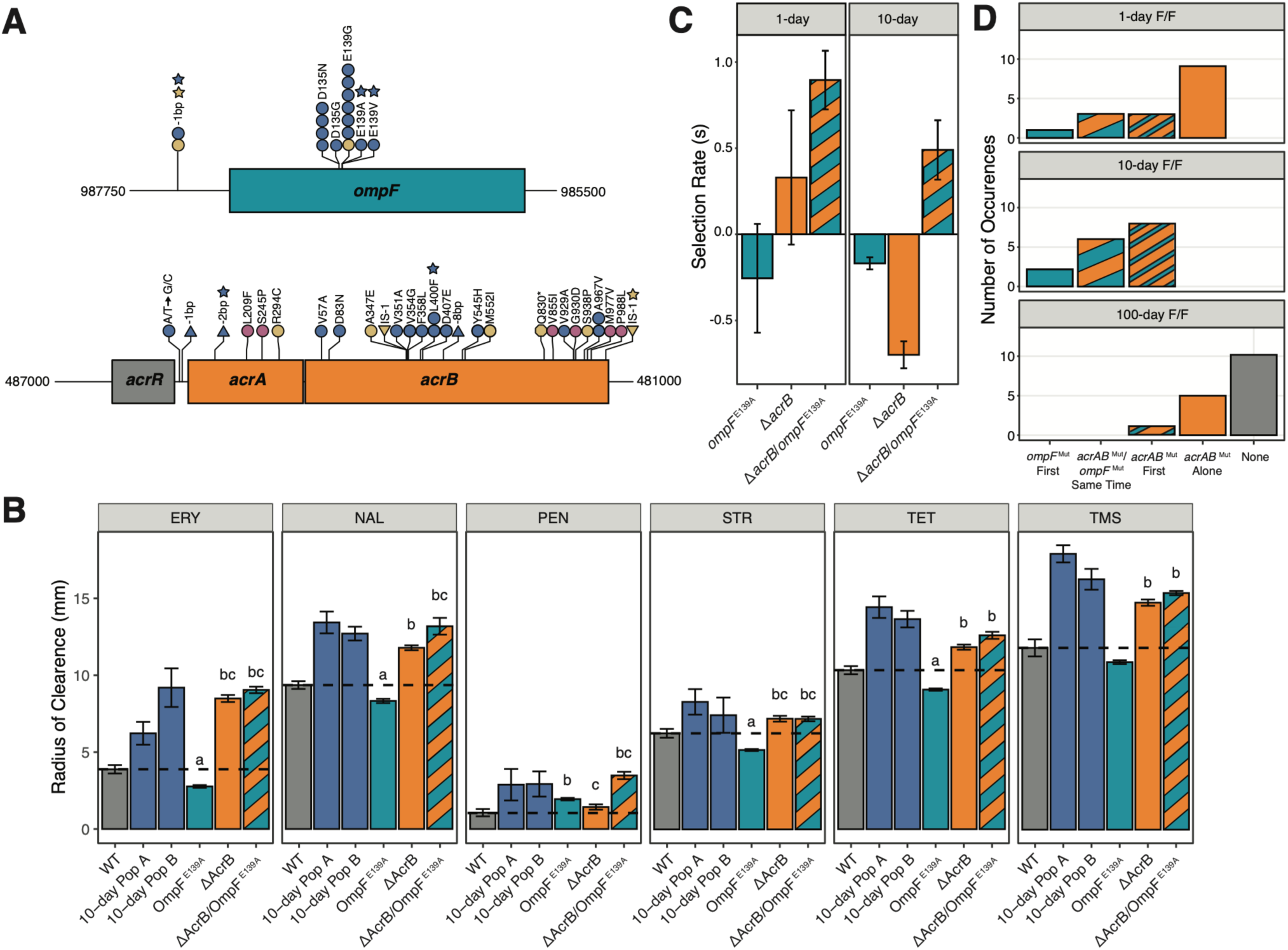
Location and effects of mutations in *ompF* and/or *acrB*. **A)** Genetic map of *ompF*, *acrR,* and the *acrAB* operon, with notable mutations annotated. Circles correspond to SNPs, upwards triangles correspond to insertions, and downward triangles correspond to deletions. Stars indicate mutations identified in the focus populations phenotyped in this study (Fig. 1C). Each circle or triangle represents one clone with an identified mutation, and its color indicates its feast/famine treatment: 1-day (yellow), 10-day (blue), 100-day (pink). **B)** Radii of clearance representing ZOI’s for the WT ancestor, clones from 10-day populations after 300 days, and reconstructed mutants for each antimicrobial. Black dashed lines indicate the WT ancestor’s radius of clearance for each antimicrobial. Letters denote significance (*p* < 0.05) (a) more resistant than the WT ancestor, (b) more susceptible than the WT ancestor, and (c) (p > 0.05) no significant difference in susceptibility relative to clones from 10-day feast/famine populations. **C)** Selection rates (*s*) of *ompF* ^E139A^*, ΔacrB*, and *ΔacrB/ompF* ^E139A^ mutants relative to the WT ancestor after 1 day and 10days of co-culture. Error bars represent +/- s.e.m. **D)** Frequency and mutational order of *ompF* and *acrB* mutations in the LTRS across the different feast/famine conditions.

To infer the likely functional consequences of mutations in *acrAB* and *ompF* on the physiology of *E. coli*, we mapped evolved nonsynonymous mutations onto the corresponding protein structures (AcrB, PDB: 1IWG; OmpF, PDB: 2OMF). In AcrB, nonsynonymous mutations clustered primarily in three critical regions: the N-terminal porter domain 1 (PN1), transmembrane helix 2 (TM2), and transmembrane helix 4 (TM4) (**Figure S1**) (Murakami et al., 2002). As TM4, and particularly the Asp407 residue, is critical for proton translocation (Su et al., 2006), TM2 structurally supports TM4 while aiding in substrate translocation (Jewel et al., 2020), and several other mutations in *acrAB* confer frame-shifts, we suspect that selection is for a general loss or reduction of AcrAB-TolC efflux pump activity . Alternatively, in OmpF, the pattern suggests selection for a more specific alteration of porin function as mutations are largely clustered around two key residues, D135 and E139. Projection of these mutations onto the structure of OmpF reveals that D135 and E139 reside within the porin channel on a key region of loop L3 associated with the constriction zone (also referred to as anti-loop 3; **Figure S2**) (Bredin et al., 2002). Mutations at D135 and E139 alter the charge of these residues from negative to neutral, likely affecting the permeability of OmpF. Thus, we hypothesize that mutations in OmpF’s constriction zone may increase access to limiting resources during feast-famine cycles, whereas reduction of AcrAB-TolC efflux activity may limit promiscuous efflux of those resources.

### Reconstructed ΔacrB and ompF ^E139A^ mutations reveal epistatic effects on susceptibility and fitness

As these genomic and structural patterns suggest that mutations in *acrAB* and *ompF* should jointly affect antimicrobial susceptibility and fitness under feast–famine conditions, we reconstructed representative mutations in the ancestral WT background and assessed their effects (Swings et al., 2018). To this end, we generated single-mutant Δ*acrB* and *ompF* ^E139A^ strains, as well as a Δ*acrB/ompF* ^E139A^ double-mutant strain. We then repeated disk-diffusion assays to determine whether these mutations recapitulated the antimicrobial susceptibility phenotypes observed in clones from 10-day feast–famine populations (Hudzicki, J., 2009). The Δ*acrB* and *ompF* ^E139A^ single-mutants produced distinct antimicrobial susceptibility profiles, and their effects varied by drug (**Figure 3B**; **Table 2**). The Δ*acrB* single mutant was more susceptible than WT to all antimicrobials tested and exhibited comparable susceptibility as clones from 10-day populations for erythromycin, penicillin and streptomycin. Alternatively, the *ompF* ^E139A^ single mutant exhibited a greater susceptibility to penicillin compared to WT, while conferring slightly increased resistance to erythromycin, nalidixic acid, streptomycin, and tetracycline. Despite the divergent effects of the single mutants, the Δ*acrB/ompF* ^E139A^ double mutant closely resembled the 10-day clones for four antimicrobials: erythromycin, nalidixic acid, penicillin, and streptomycin. These results support a contribution of both *acrB* and *ompF* to the evolved antimicrobial susceptibility phenotype, while also revealing positive epistasis between the two mutations as the increase in susceptibility in the double-mutant is greater than the additive effect from the two single-mutants.

**Table 2:** Reconstructed *ΔacrB and ompF* ^E139A^ mutations have distinct effects on antimicrobial susceptibility.

| | 10-day F/F | $ompF^{E139A}$ | | $\Delta acrB$ | | $\Delta acrB/ompF^{E139A}$ | |
| --- | --- | --- | --- | --- | --- | --- | --- |
| Antimicrobial | ROI (mm) | ROI (mm) | p-value (adj.) | ROI (mm) | p-value (adj.) | ROI (mm) | p-value (adj.) |
| ERY | 7.59 | 2.72 | 0.0035 | **8.35<sup>^</sup>** | **0.38** | **8.9<sup>^</sup>** | **0.28** |
| NAL | 12.9 | 8.2 | 2.00E-07 | 11.6 | 0.024 | **13<sup>^</sup>** | **0.87** |
| PEN | 2.86 | **1.91<sup>^</sup>** | **0.31** | **1.41<sup>^</sup>** | **0.12** | **3.43<sup>^</sup>** | **0.4** |
| STR | 7.73 | 5.08 | 0.0095 | **7.08<sup>^</sup>** | **0.71** | **7.07<sup>^</sup>** | **0.71** |
| TET | 13.8 | 8.93 | 5.30E-07 | 11.7 | 6.60E-04 | 12.4 | 0.012 |
| TMS | 16.8 | 10.7 | 9.90E-08 | 14.5 | 1.10E-03 | 15.1 | 5.10E-03 |

ROI (mm) = Radius of Inhibition
<sup>^</sup>Reconstructs susceptibility observed in 10-day F/F populations

We next asked whether these mutations also affected fitness under feast–famine conditions. Here, we competed each reconstructed mutant against the WT ancestor under 1-day and 10-day culture conditions, using selection rate (*s*) as a proxy for relative fitness (**Figure 3C**). The *ompF* ^E139A^ single mutant exhibited a negative selection rate in both the 1-day and 10-day culture conditions, indicating that this substitution is deleterious on its own at both timepoints (*s_ompF_*_, 1-day_ = -0.255; *s_ompF_*_, 10-day_ = -0.169). In contrast, the Δ*acrB* single mutant showed condition-dependent fitness effects. Under 1-day conditions, the Δ*acrB* mutation is beneficial but deleterious under 10-day conditions (*s_acrB_*_, 1-day_ = 0.330; *s_acrB_*_, 10-day_ = -0.700), suggesting a fitness trade-off for Δ*acrB* between growth and survival. Strikingly, the Δ*acrB/ompF* ^E139A^ double mutant outcompeted the WT ancestor in both 1- and 10-day conditions, indicating a substantial advantage compared to either single mutant (s*_acrB/ompF_*_, 1-day_ = 0.895; s*_acrB/ompF_*_, 10-day_ = 0.490). Moreover, the selection rates of the Δ*acrB/ompF* ^E139A^ double mutant differed greatly from the expected additive fitness phenotype in both conditions, consistent with positive epistasis between *ompF* ^E139A^ and Δ*acrB* under 1-day conditions (ℇ = 0.82), and reciprocal sign epistasis under 10-day conditions (ℇ = 1.359).

Given that *ompF* ^E139A^ is only beneficial in the presence of Δ*acrB,* we investigated the order in which mutations in *acrAB* and *ompF* arose across LTRS populations during the full 900 days of experimental evolution and found that the evolutionary trajectories are consistent with the fitness of the reconstructed mutants (**Figure 3D; Supplemental Dataset S2**). Under 1-day feast/famine conditions, populations overwhelmingly either evolve mutations solely in *acrAB* (9/16 populations). Alternatively, under 10-day feast/famine conditions, populations most frequently evolve mutations in both *acrAB* and *ompF* (14/16 populations). In these populations, mutations either arise in *acrAB* first (8 populations) or in both *acrAB* and *ompF* within the same sequencing timepoint (6 populations). Two exceptions were the 10-day populations, P409 and P418, where only an *ompF* mutation was initially detected. However, closer inspection revealed that these populations instead evolved mutations in *acrD,* a homolog to *acrB* which also forms an efflux pump complex with AcrA and TolC. Thus, even this exception is consistent with a broader pattern in which altered porin function evolves alongside disruption of AcrAB-like efflux activity. As such, when combined with the fitness of the reconstructed mutants, it appears that mutations in *ompF* are compensatory and provide benefits that overcome the deleterious effects of *acrAB* mutations under the 10-day starvation conditions. Lastly, populations evolving under 100-day feast/famine conditions generally only evolve mutations in *acrAB* or no mutations in either the *acrAB* or o*mpF* loci. This suggests that the selective pressures experienced under 100-day feast/famine conditions are distinct from 10-day feast/famine conditions, and that different genotypes are favored when navigating extreme resource fluctuations.

### ompF ^E139A^ compensates for reduced efflux by ΔacrB during 10-day culture

Initially, we hypothesized that substitutions in the OmpF constriction loop would enable greater uptake of resources under feast/famine conditions. However, given that the *ompF* ^E139A^ mutant confers increased antimicrobial resistance to several drugs, it’s more likely that substitutions in the constriction loop instead reduce uptake. To better dissect how mutations in *acrB* and *ompF* influence *E. coli* physiology, we first assessed their growth in cell-free conditioned LB medium. Here, we conditioned the medium by growing the WT ancestor for 24 h before filtering out the cells, providing a relevant resource-depleted medium for competitive fitness assays (**Figure 4A**). In contrast to fitness in fresh LB medium, the Δ*acrB* mutation is deleterious (*s* = -0.332), and the *ompF* ^E139A^ mutation is beneficial in depleted LB media (*s* = 0.460). As disruption of AcrA or AcrB should lead to reduced efflux and retention of metabolites, this may prove deleterious if toxic metabolic byproducts begin to accumulate in the medium after 24 h of growth. Consistent with this, the Δ*acrB/ompF* ^E139A^ double mutant is weakly beneficial (*s* = 0.178), further supporting our revised hypothesis that the *ompF* ^E139A^ mutation reduces OmpF import activity and compensates for the disruption of AcrAB-TolC function.

**Figure 4:**
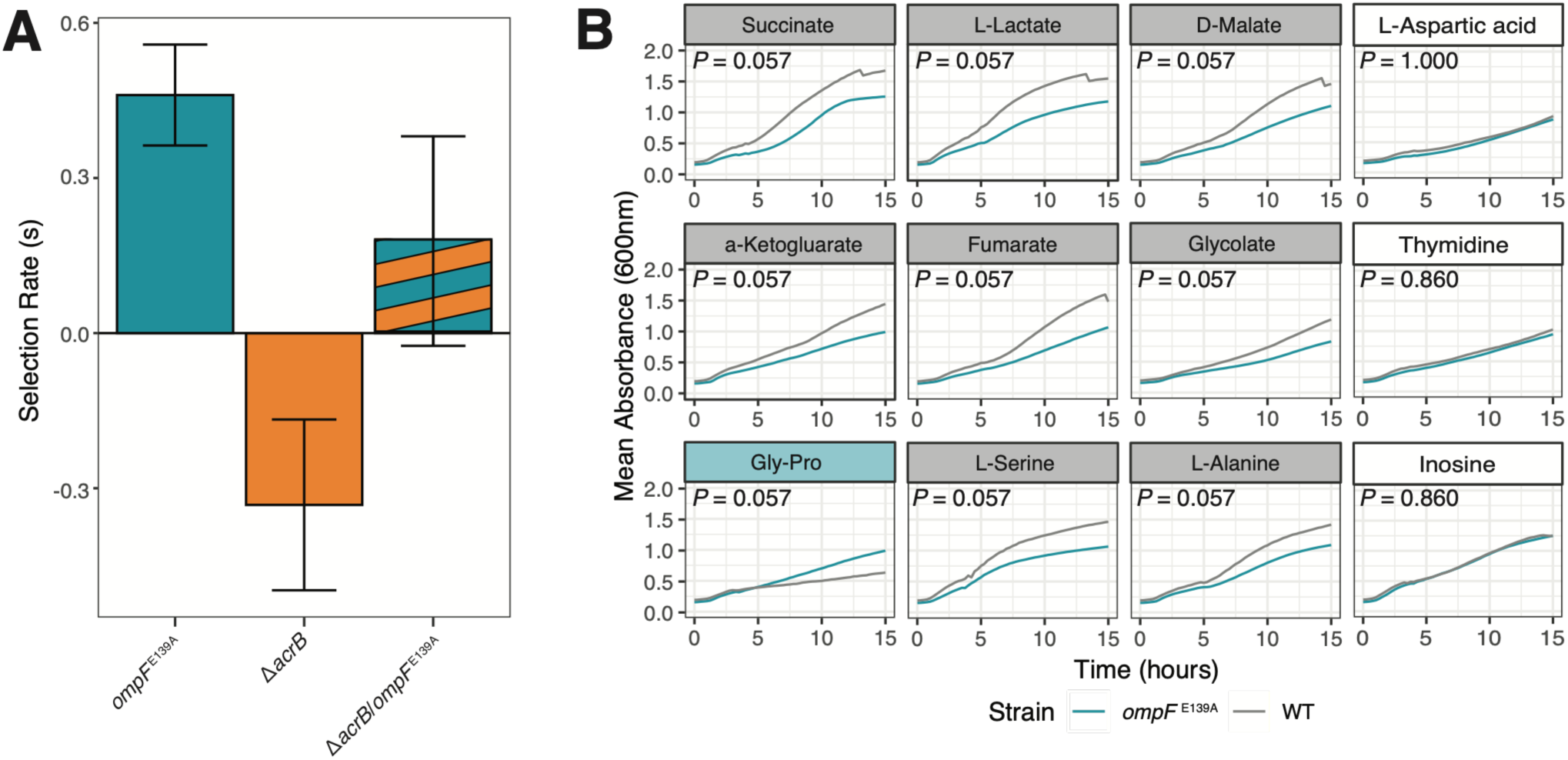
Patterns of fitness in depleted LB media and growth across various carbon resources suggest reduced import by *ompF* ^E139A^. **A)** Selection rates (*s*) of *ompF* ^E139A^*, ΔacrB*, and *ΔacrB/ompF* ^E139A^ mutants relative to the WT ancestor after 24 hours of culture in cell-free WT conditioned media. Error bars represent +/- s.e.m. **B)** Selected plots illustrating growth profiles for *ompF* ^E139A^ and the WT ancestor across single carbon sources. Colored panels indicate the strain with significantly greater AUC_e_: *ompF* ^E139A^ (cyan), WT (grey), none (white). For all growth profiles and AUC_e_ measurements, see Supplemental Figures S3 and S4.

To test whether *ompF* ^E139A^ broadly restricts resource uptake, we compared growth of the *ompF* ^E139A^ single mutant and WT ancestor across 65 carbon sources (**Figure 4B; Figure S3-S4; Supplemental Dataset S3**). Consistent with reduced import, the *ompF* ^E139A^ mutant showed lower growth than WT on nearly half of the tested resources. Specifically, *ompF* ^E139A^ produced significantly lower area under the curve (AUC_e_) values on 30 carbon sources (∼46% of resources) and outperformed WT on only three carbon sources (∼4.6% of resources). Among these resources, *ompF* ^E139A^ performed worst on fumarate, lactate, and succinate (∼ 35% decrease in AUC_e_), and best on glycine dipeptides and glyoxylic acid (36-63% increase in AUC_e_). Together, these results suggest that *ompF* ^E139A^ reshapes membrane permeability, compensating for the fitness cost of impaired AcrAB-TolC efflux during 10-day culture.

## Discussion

Previously, we demonstrated that adaptation to fluctuating nutrient availability selects for mutations that confer trade-ups, optimizing growth and survival across repeated cycles of feast and famine (Behringer et al., 2024). Here, we show that this process can also generate collateral changes in antimicrobial susceptibility despite the complete absence of antibiotic selection. Specifically, populations evolving under intermediate (10-day) feast-famine cycles repeatedly evolved increased antimicrobial susceptibility to multiple drug classes, a latent phenotype arising as an indirect consequence of adaptation to fluctuating resources. By combining longitudinal population genomics with allele reconstruction and competitive fitness assays, we identified mutations affecting the multidrug efflux pump AcrAB-TolC and the outer membrane porin OmpF as major contributors to this phenotype. Importantly, the fitness effects of these mutations depend on both ecological context and genetic background, illustrating how the adaptive trajectories of populations evolving under fluctuating conditions can be shaped by environment-dependent epistasis. Together, these findings reveal how fluctuating environments shape the genetic architecture of adaptation through environment-dependent epistasis, producing historically contingent evolutionary trajectories and latent phenotypes.

Our results suggest that the repeated evolution of mutations in *acrAB* and *ompF* reflects selection on balancing import and efflux under fluctuating resource conditions. AcrAB-TolC exports a broad range of substrates, including metabolic intermediates, fatty acids, bile salts, detergents, and antibiotics (Elkins & Nikaido, 2002; Ramaswamy et al., 2017; Yu et al., 2003). In contrast, OmpF provides a major route for the passive import of small hydrophilic molecules into the cell, nutrients and antimicrobials alike (Nikaido, 1992). The repeated evolution of *ompF* is also consistent with the Self-Preservation and Nutritional Competence (SPANC) framework, which predicts that adaptation to prolonged nutrient limitation frequently involves changes in outer membrane permeability and nutrient acquisition (Ferenci, 2005). While we initially hypothesized that mutations in the OmpF constriction loop would increase nutrient uptake during prolonged starvation, our results instead indicate that this mutation more likely reduces import, accounting for the slightly increased resistance and reduced growth on single carbon resources exhibited by the *ompF*^E139A^ mutant. Previous work demonstrated that disruption of AcrB (D408A) promotes intracellular retention of Vitamin D and oxidized fatty acids in LB medium (Wang-Kan et al., 2021), suggesting that reduced efflux may provide an advantage by conserving biosynthetically expensive metabolites during periods of rapid growth. However, prolonged retention of these metabolites, particularly short-chain fatty acids, may become detrimental during extended stationary phase as their accumulation can become toxic (Royce et al., 2013). Together, these observations support a working model in which the alteration of OmpF permeability partially restores physiological homeostasis by limiting further influx of extracellular metabolites, thereby compensating for impaired efflux while simultaneously increasing susceptibility to antimicrobial compounds. Thus, mutations in *acrB* and *ompF* appear to reflect optimization of import and efflux under fluctuating nutrient conditions, consistent with observations of gram-negative bacteria prioritizing *ompC* in cells missing TolC transporters (Dastidar et al., 2007; Misra & Reeves, 1987; Morona & Reeves, 1982).

Selection rates for the Δ*acrB* and *ompF* ^E139A^ mutations further demonstrate that the adaptive value of these variants depends strongly on ecological context. The fitness effects of the reconstructed alleles differ across one-day and ten-day culture conditions, showcasing how the selective landscape experienced by the population changes continuously as cultures progress from exponential growth through stationary phase and into prolonged starvation. Rather than occupying a single static fitness landscape, evolving populations instead traverse a dynamic fitness seascape in which the selective value of individual mutations changes over time (Maltas et al., 2021; Mustonen & Lässig, 2009; Stone & Behringer, 2026; Trubenová et al., 2019). Loss of AcrAB function is beneficial during the first day of growth yet becomes deleterious during prolonged starvation. Addition of the *ompF* ^E139A^ mutation reverses this fitness cost, producing a double mutant that outcompetes the ancestral strain under both ecological conditions. This provides an experimental example of how evolved mutations that are individually deleterious during one phase of the feast-famine cycle can become beneficial when combined with subsequent mutations that restore physiological balance. These findings illustrate how temporal variation in selection can reshape epistatic interactions, resulting in adaptive trajectories that would not be predicted from measurements made at a single point in the growth cycle.

The repeated order in which these mutations arose across independently evolving populations further suggests an important role for historical contingency. Mutations affecting *acrAB* consistently appeared before mutations in *ompF*, mirroring the fitness landscape inferred from the reconstructed mutants. Because the *ompF* ^E139A^ mutation is deleterious in the ancestral background but beneficial following disruption of *acrB*, its accessibility depends upon prior evolutionary history. Historical contingency arises when genetic interactions alter the accessibility of future evolutionary trajectories, often through sign-epistasis (Blount et al., 2018). For instance, an A258T substitution in GltA, which increases citrate synthase activity, created the physiological conditions that permitted the evolution of aerobic citrate utilization in the LTEE (Blount et al., 2008, 2012; Quandt et al., 2015). In *Pseudomonas*, epistatic interactions between chromosomal mutations permit the carriage of multidrug resistance plasmids (Loftie-Eaton et al., 2017). While in nature, sign-epistasis underlies the evolved changes between the 5S rRNA genes of *Vibrio alginolyticus* and *Vibrio proteolyticus* (Weinreich et al., 2005).

Our results extend this concept by showing that dynamic ecological conditions can contribute to historical contingency by altering the fitness effects of individual mutations through time. In this system, the transition from nutrient abundance to prolonged starvation creates conditions under which loss of AcrAB function first becomes advantageous and subsequently permits compensatory evolution in OmpF. Thus, historical contingency emerges not only through epistatic interactions among mutations but also through the temporal structure of the selective environment itself (**Figure 5**).

**Figure 5.**
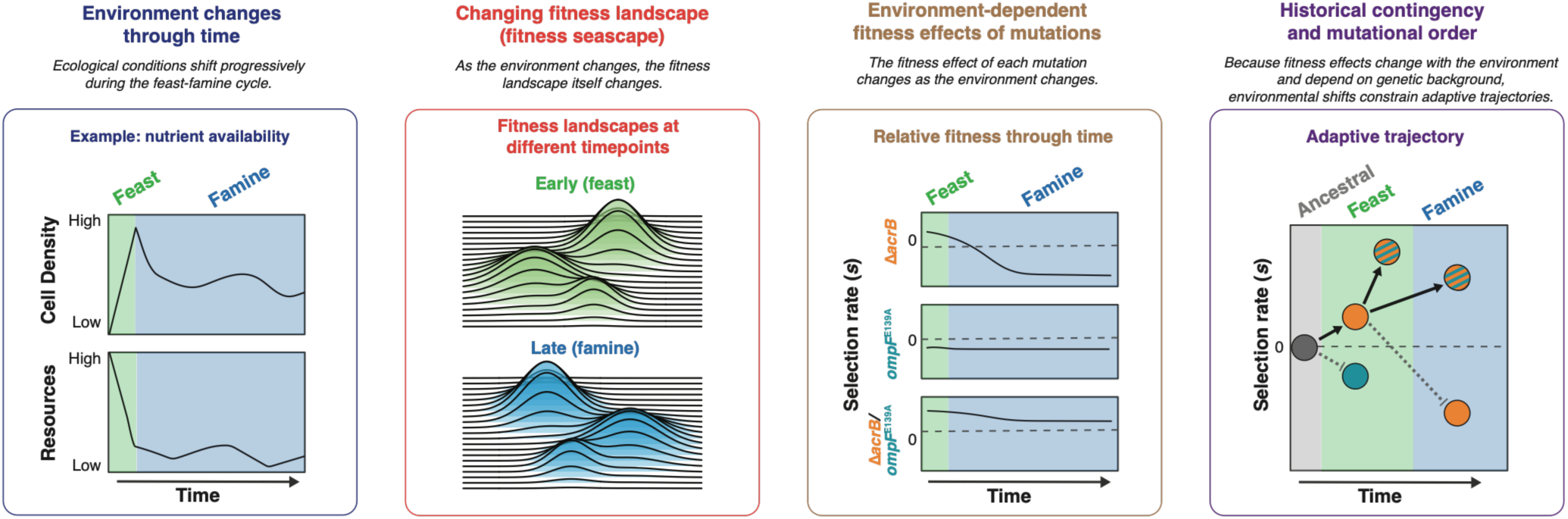
Conceptual model illustrating how fluctuating nutrient availability generates environment-dependent epistasis and historical contingency. During repeated feast-famine cycles, feedbacks between nutrient availability and cell density continuously reshape the selective environment experienced by the population through time (left). As ecological conditions shift, the fitness landscape itself changes, creating a dynamic fitness seascape and modifying the fitness effects of individual genotypes across the growth cycle (center left). Consequently, the fitness effects of *ΔacrB, ompF* ^E139A^, and the *ΔacrB/ompF* ^E139A^ double mutation depend on environmental context, resulting in environment-dependent epistasis (center right). These changing fitness effects constrain the accessibility of adaptive mutations, favoring a sequential evolutionary trajectory in which initially beneficial mutations facilitate subsequent mutations that together produce a net fitness advantage across both feast and famine (right).

Our findings extend a growing body of work showing that pleiotropic effects of adaptive mutations produce latent phenotypes (Knöppel et al., 2017; Lamrabet et al., 2019; Paaby & Rockman, 2013; Payne & Wagner, 2014; Rendueles & Velicer, 2020). In the LTEE, reduced expression of the maltose transporter *lamB* during adaptation to glucose-rich conditions incidentally resulted in resistance to bacteriophage λ (Meyer et al., 2010). Likewise, reduction of stress-associated gene expression via mutations in RpoB during adaptation to thermal stress incidentally results in rifampicin resistance due to alterations in the antibiotic binding target (Rodríguez-Verdugo et al., 2013). Our results extend these observations by demonstrating that adaptation to fluctuating nutrient availability can result in increased broad-range antimicrobial susceptibility through the inherent pleiotropic trade-offs associated with broad-range metabolite transport. These findings reinforce the view that increased antimicrobial sensitivity is not solely a consequence of relaxed selection, or negative selection on a costly trait, when antibiotic pressure is removed (Burmeister et al., 2020; Lamrabet et al., 2019; Maltas et al., 2020). Instead, increased antimicrobial susceptibility may also emerge as a physiological consequence of adaptation to resource limitation, a challenge regularly encountered by microbes.

In summary, our results demonstrate that repeated feast-famine cycles produce predictable evolutionary trajectories in which adaptation to fluctuating resource availability generates antimicrobial susceptibility as a latent phenotype. By linking parallel genomic evolution with allele reconstruction and ecological fitness measurements, we show that environment-dependent epistasis and historical contingency emerge from the changing selective pressures experienced across the microbial growth cycle. More broadly, these findings provide additional evidence that ecological history can reshape the genetic architecture of adaptation, with important consequences for antimicrobial susceptibility.

Because fluctuating resource availability is ubiquitous in natural, host-associated, and laboratory environments, the temporal structure of resource availability is likely to be an important determinant of both evolutionary trajectories and the latent phenotypes they produce.

## Acknowledgements

We would like to thank C.H.J., J.C.M, R.S., and A.S.W. for their early comments and contributions to this work. Genomic sequencing services were provided by Indiana University Center for Genomics and Bioinformatics. High-performance computing resources were provided by Vanderbilt University’s Advanced Computing Center for Research and Education (ACCRE).

## Study Funding

This work was supported by the National Institute of General Medical Sciences Grant R35GM150625 (M.G.B.) and supplement to support S.B.W. Additional funds were provided by the Vanderbilt Evolutionary Studies Initiative (A.G.B.; Z.C.) and the Vanderbilt Undergraduate Summer Research Program (VUSRP) (N.F.).

## Data Availability

Raw sequencing reads associated with this project are available through NCBI’s Sequencing Read Archive (BioProject Numbers: PRJNA1502908 and PRJNA532905) All R code and raw data used to generate figures for this manuscript are available at the Behringer Lab’s GitHub repository: https://github.com/BehringerLab/Environment_Dependent_Epistasis.

## Conflict of Interest

The authors report no conflicts of interest.

---

**Supplemental Figure S1:**
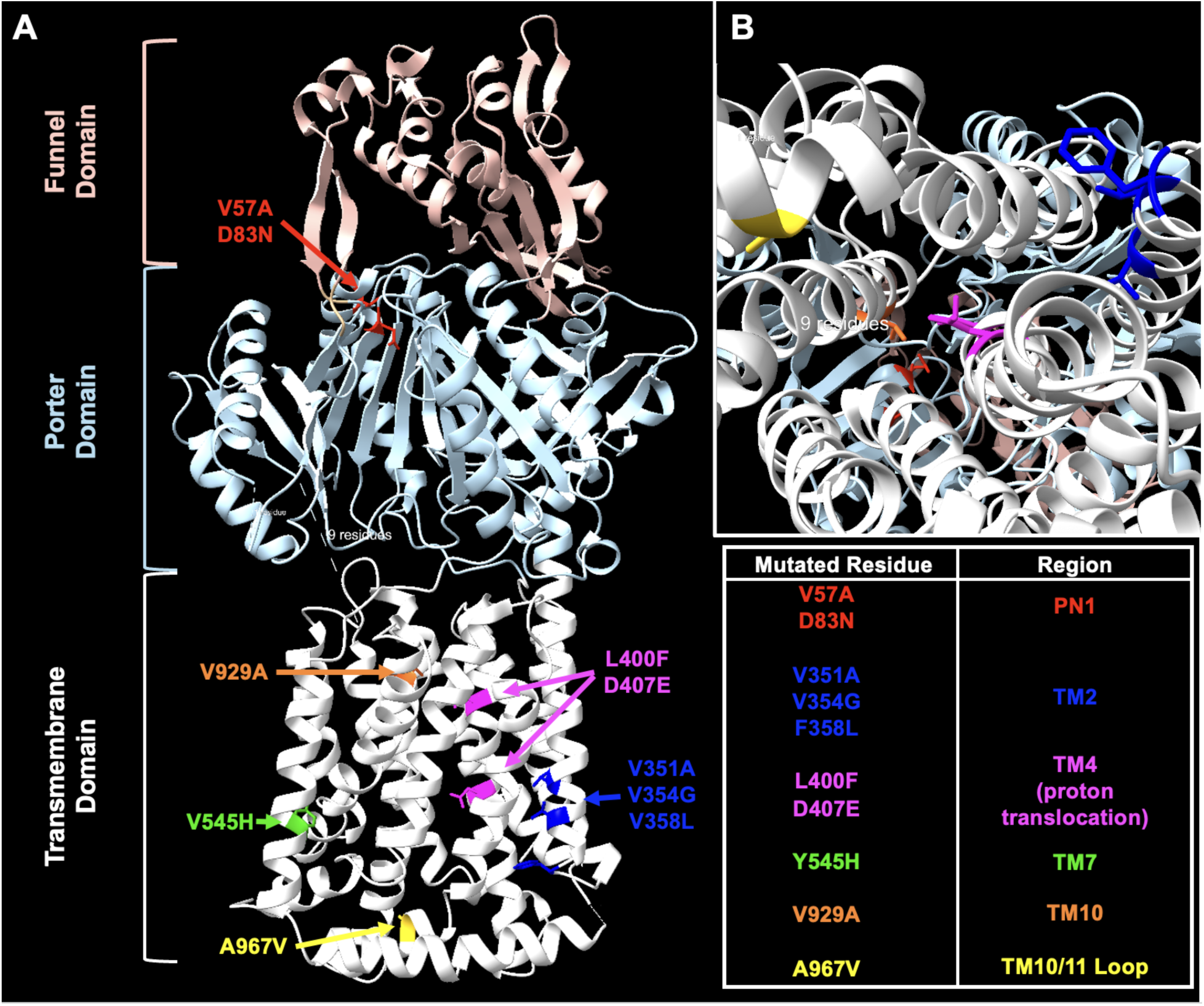
Structure of AcrB and location of nonsynonymous variants. Structure based on 1IWG. **A)** View is from the side of AcrB, where AcrB is visualized as colored ribbons based on domain: transmembrane (white), porter (light blue), and funnel (light pink). Nonsynonymous variants are visualized as atoms and highlighted in different colors based on their location in the protein (see the inset table). **B)** View is looking into the protein from the periplasm to visualize the proton translocation channel. This figure was made with Chimera X.

**Supplemental Figure S2:**
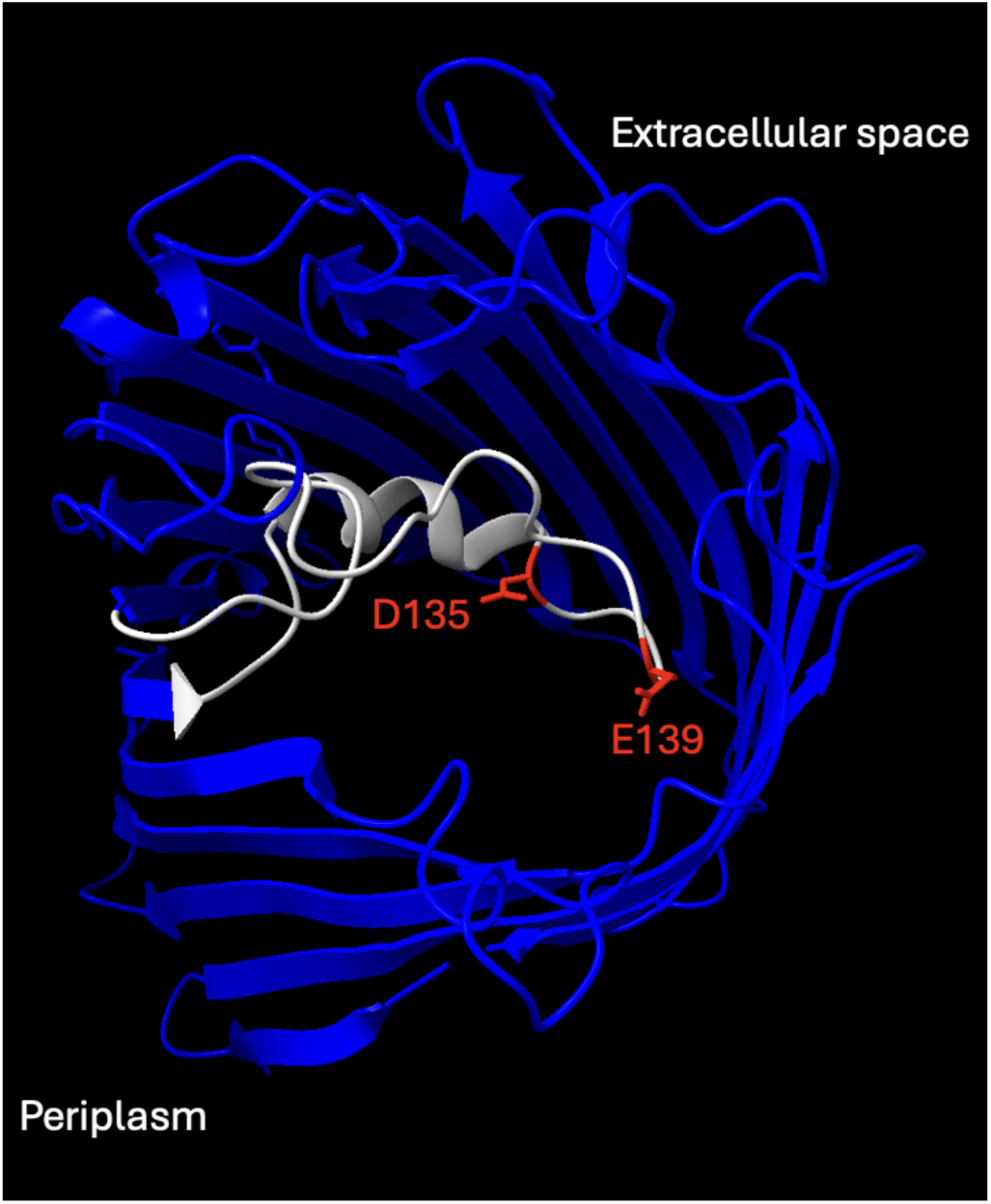
Structure of OmpF and location of D135 and E139. Structure based on 2OMF. View is from the extracellular side, looking through the porin to the periplasm. OmpF is visualized as blue ribbons, and the constriction zone (Loop 3, L3) is highlighted in white from residues R122 - G157. D135 and E139 are shown as atoms and highlighted in red. This figure was made with Chimera X.

**Supplemental Figure S3:**
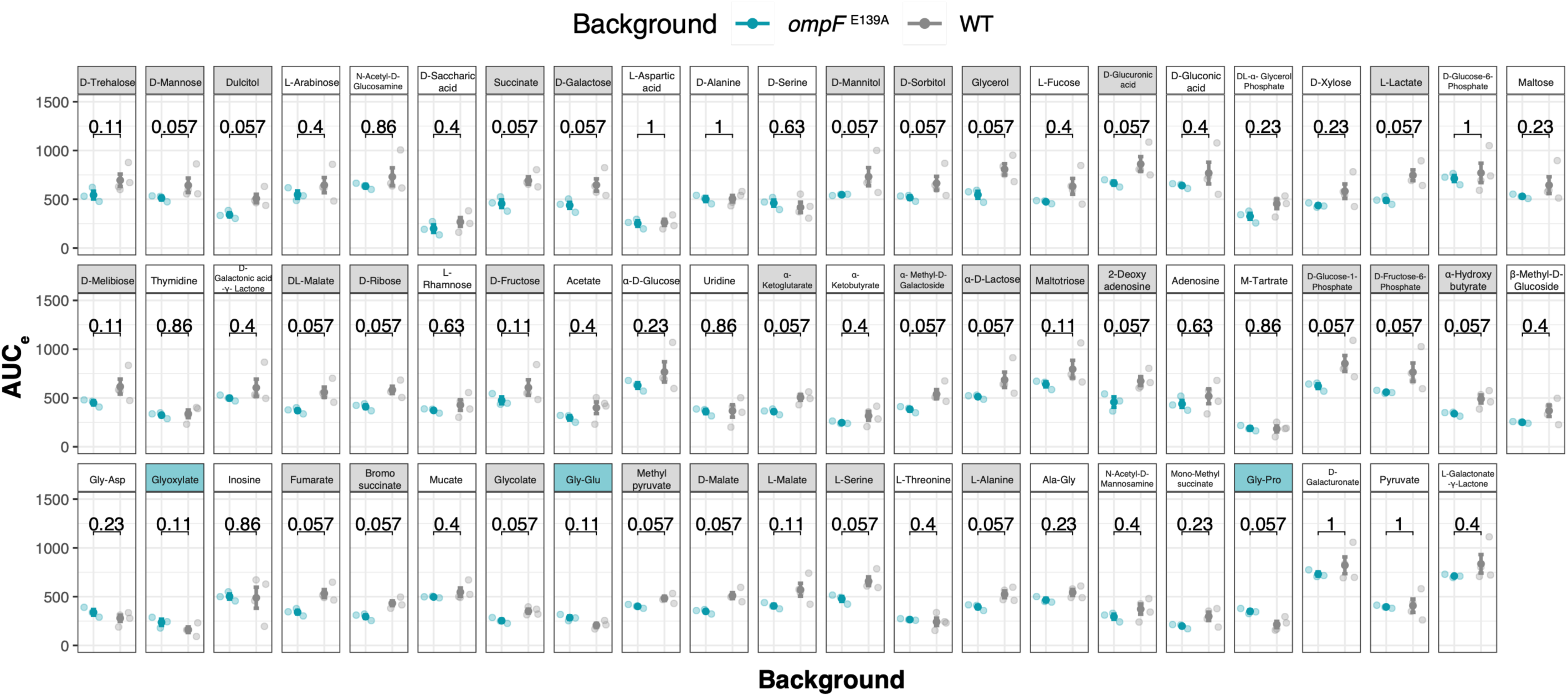
Comparison of WT and the *ompF* ^E135A^ mutant strain growth across 65 carbon sources. Growth is quantified as the empirical area under the curve (AUCe). Each resource is highlighted based on the strain that exhibited the best growth on the carbon resource: *ompF* ^E139A^ (teal), WT (gray), or none (white). Individual AUC_e_ values are indicated by transparent circles, mean AUC_e_ values are indicated by opaque circles, and error bars illustrate +/- s.e.m. AUC_e_ was compared for each resource using a pairwise t-test. P-values are shown for each resource.

**Supplemental Figure S4:**
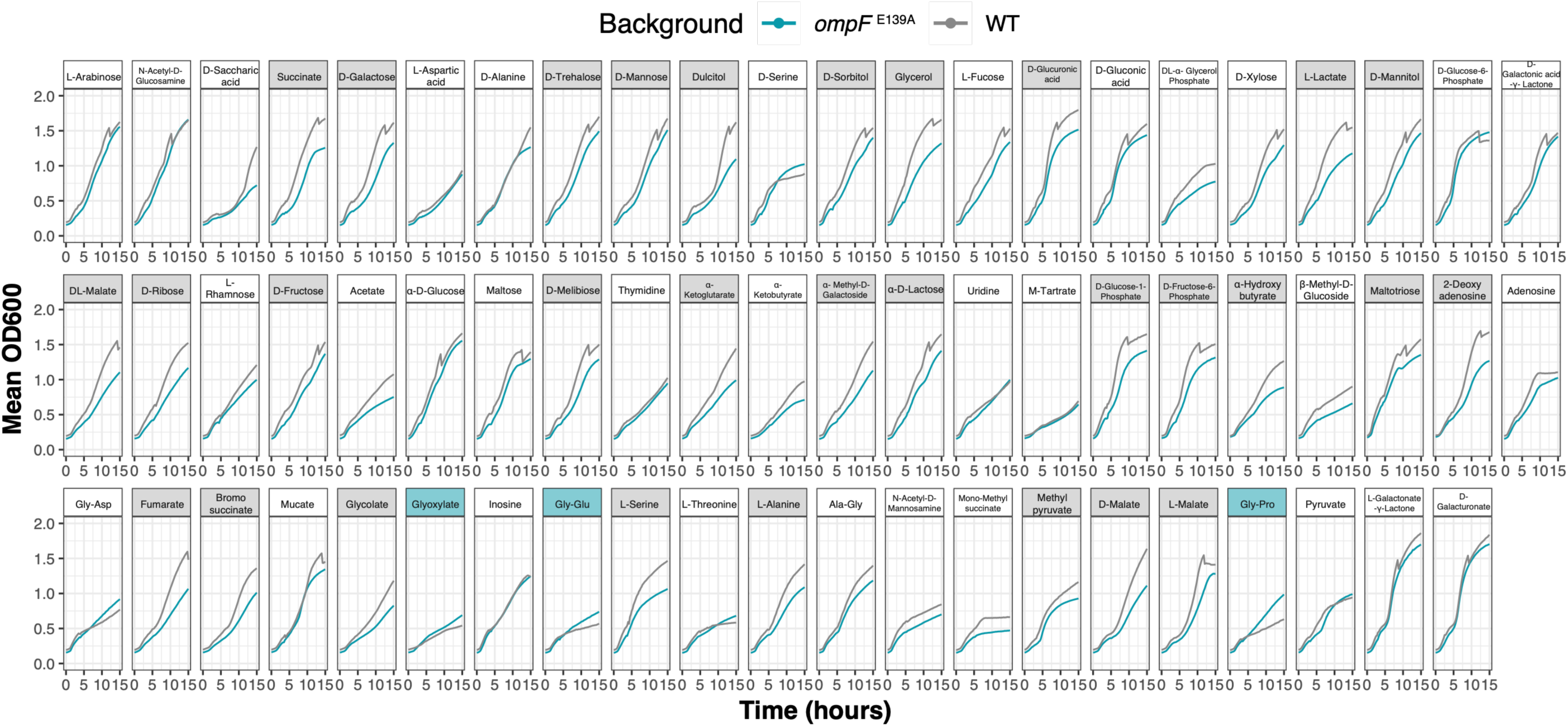
Growth curves for WT and the *ompF* ^E139A^ mutant strain across 65 carbon sources. Growth is visualized as mean growth across three replicates for each strain. Each resource is highlighted based on the strain that exhibited the best growth on the carbon resource: *ompF*^E139A^ (teal), WT (gray), or none (white) (see **Figure S3**).

**Supplemental Dataset 1:** Mutation profiles for evolved clones assessed in this study.

**Supplemental Dataset 2:** Mutations in *ompF* or *acr* loci across all sequenced timepoints in the LTRS.

**Supplemental Dataset 3:** Growth profiles of *ompF*^E139A^ and WT across 65 carbon sources

## Notes

https://github.com/BehringerLab/Environment_Dependent_Epistasis

### Competing Interest Statement

The authors have declared no competing interest.

